# *Takakia* possesses a key marker of embryophyte sporopollenin

**DOI:** 10.1101/2024.01.17.576121

**Authors:** Dae-Yeon Suh, Damanpreet K Sraan, Neil W Ashton

**Affiliations:** Department of Chemistry and Biochemistry, University of Regina, Canada; Department of Biology, University of Regina, Canada

**Author notes:** Correspondence to: Dae-Yeon Suh, Department of Chemistry and Biochemistry, University of Regina, Regina, Saskatchewan, S4S 0A2, Canada., Correspondence to: Neil W Ashton, Department of Biology, University of Regina, Regina, Saskatchewan, S4S 0A2, Canada.

**Keywords:** Anther-Specific Chalcone synthase-Like, Sporopollenin, *Takakia*

## Abstract

The enigmatic moss, *Takakia lepidozioides*, possesses a particular type III polyketide synthase, ASCL (Anther-Specific Chalcone synthase-Like), that is an identifying marker for genuine sporopollenin in the walls of embryophyte spores and pollen grains. By contrast, a survey of all algae with sequenced genomes confirms that they do not possess ASCL and, therefore, their spore walls are not composed of sporopollenin.

## Introduction

Sporopollenin (SP) is the chemically resistant wall material of plant spores and pollen, which provides protection from subaerial stresses. For most of the time since the term was coined by Zetzsche in 1931 (Zetzsche and Kälin, 1931; Zetzsche and Vicari, 1931) resistance to acetolysis has been the sole criterion for the identification of SP. This chemically imprecise definition has resulted in numerous claims for the existence of SP or SP-like material in algae and various microorganisms (Suh and Ashton 2022 and references therein). Based on more recent chemical analyses (reviewed by Grienenberger and Quilichini 2021), the current view of SP is that it is a polymer composed of polyhydroxylated polyketides, hydroxylated aromatics and fatty acid derivatives, crosslinked via ester and ether bonds and oxidative C−C coupling. A particular polyketide synthase (PKS), ASCL, plays a central role in the polyketide biosynthetic pathway and is proposed to provide hydroxylated polyketides as SP precursors (Kim et al. 2010; Colpitts et al. 2011; Suh and Ashton 2022). Following an extensive BLASTp survey of plant genomes, we discovered that with a single exception, the marine monocot, *Zostera marina*, which has exineless pollen (Olsen et al. 2016), representatives of all major embryophyte clades possess ASCL in stark contrast to algae, including Charophytes, which do not. This led us to propose a new definition for SP as follows to distinguish genuine SP from SP-like compounds such as the acetolysis-resistant algaenans possessed by some Chlorophytes and to use ASCL as an identifying marker for the presence of SP in spore and pollen grain walls (Suh and Ashton 2022).

‘Sporopollenin is a chemically resistant complex heteropolymer present in the outer walls of spores and pollen grains and is composed partly of hydroxylated polyketides derived from the conserved polyketide pathway, which involves ASCL.’

At the time we made this definition, the genome of the phylogenetically enigmatic plant, *Takakia*, of which there are only two species, *Takakia lepidozioides* and *Takakia ceratophylla*, was not available. This plant is fascinating because of its phylogenetic affinities to algae, hornworts and liverworts as well as mosses and for a long time its phylogenetic position remained uncertain. Recently, *Takakia* has been shown to be sister to all other mosses including *Sphagnum* (Liu et al. 2019) and to have diverged from the Last Common Ancestor (LCA) of embryophytes after the hornworts and liverworts (Hu et al. 2023). It shares 1118 genes, which are absent in *Physcomitrium patens*, with one or more Charophytes and 1665 genes, which are also absent in *P. patens*, with liverworts and hornworts (Hu et al. 2023). The functions of most of these genes are unknown but could be the basis for shared characteristics of *Takakia* with liverworts and algae but not mosses (Hu et al. 2023). It is presumed by Hu et al. (2023) that most of them were present in the LCA of bryophytes and were secondarily lost in *P. patens* by reductive evolution. For these reasons, we were keen to discover whether the recently sequenced *Takakia lepidozioides* genome possesses an *ASCL* gene indicating the probable presence of SP in its spore wall, which like those of other mosses comprises intine, exine and perine layers (Renzaglia et al. 1997), and also to reinforce our contention that ASCL is present in representatives of all major embryophyte groups.

## Methods

We performed BLASTp searches against the *Takakia lepidozioides* genome database (https://www.takakia.com/blast/blast_cs.html, accessed on August 31, 2023) with PpASCL and PpORS (*Physcomitrium patens* 2′-OxoalkylResorcinol Synthase), a bryophyte/charophyte-specific type III PKS, as query sequences. This yielded sixteen putative type III PKS models, which contained the catalytic Cys-His-Asn triad and signature sequences (G/A)FGPG (Suh et al. 2000). Similarly, putative algal type III PKS sequences were retrieved by BLASTp searches against each algal genome in PhycoCosm (https://mycocosm.jgi.doe.gov/algae/algae.info.html, accessed on September 1, 2023). In cases of fusion proteins, portions of sequences that matched type III PKS sequences from the same or related species were taken for further analysis. Representative embryophyte type III PKS sequences were retrieved from Phytozome 13 (https://phytozome-next.jgi.doe.gov/) as described previously (Aslam et al. 2022). The sequences used for tree reconstruction (Table 1) were aligned by the MUSCLE method in MEGA 11 (Tamura et al 2021), and a Maximum Likelihood (ML) phylogenetic tree (Fig. 1) was reconstructed in MEGA 11 using the JTT substitution model. The initial tree was created using the default NJ/BioNJ method, and tree improvement was performed using the nearest-neighbor-interchange ML heuristic method. Support for the tree was measured using 1,000 bootstrap replicates.

**Table 1.**
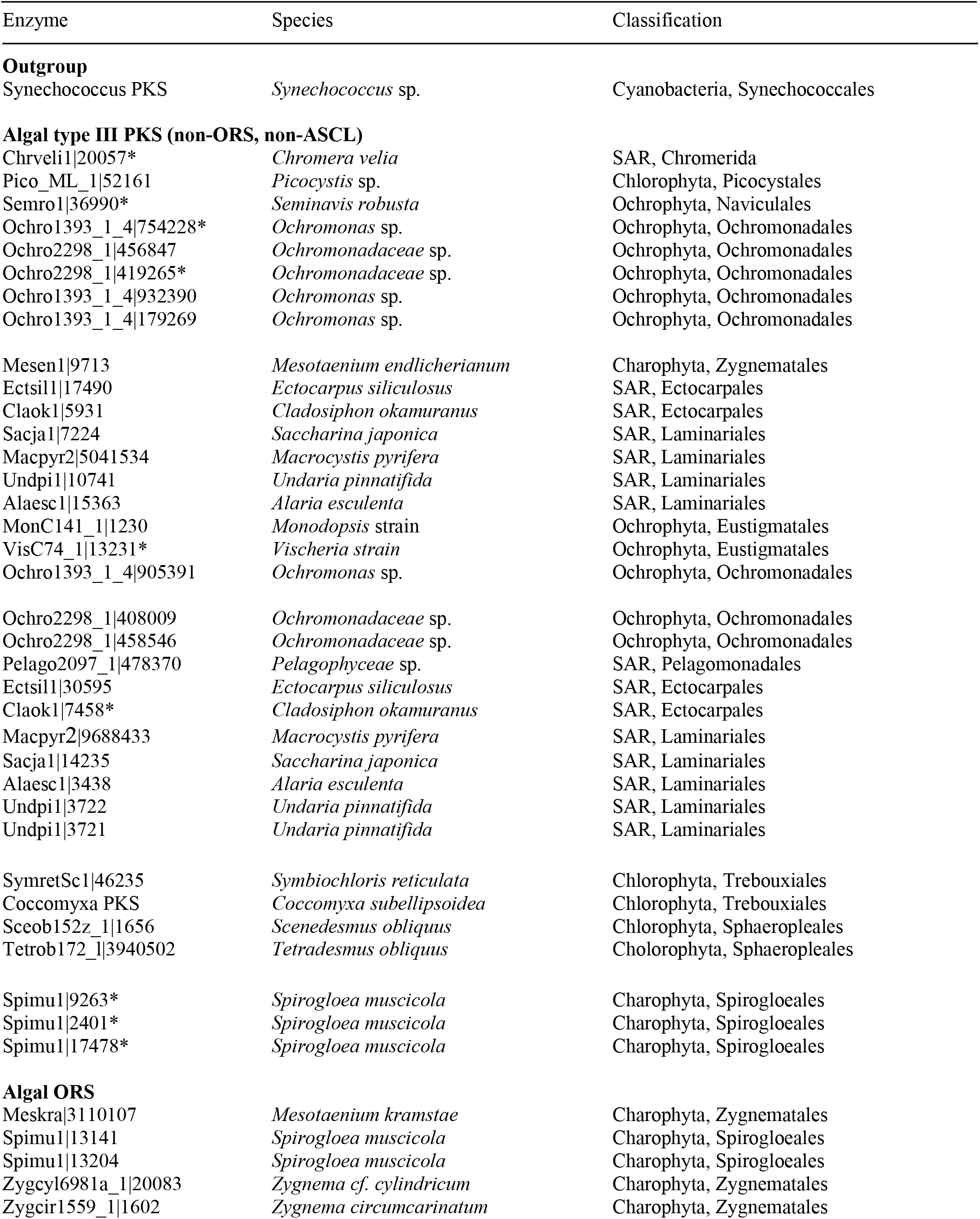
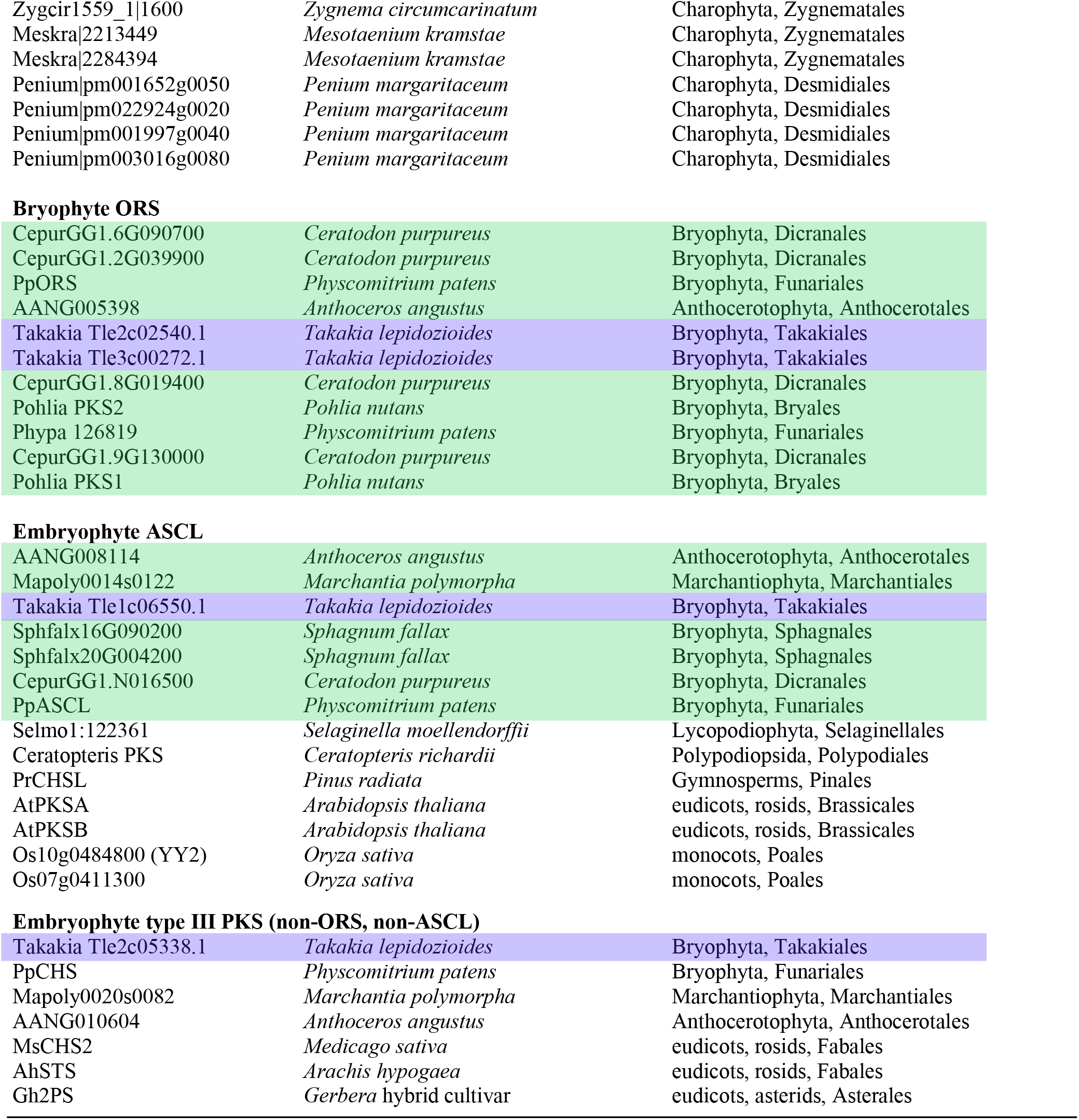
Plant type III polyketide synthases used for tree reconstruction. Type III PKSs are listed in the same order of their appearance (from the top) in the tree (Fig. 1) before collapsing some of the clades. *Takakia* enzymes and other bryophyte enzymes are highlighted in blue and green, respectively, as in Figure 1. Fusion proteins containing a type III PKS domain are indicated with asterisks.

**Figure 1.**
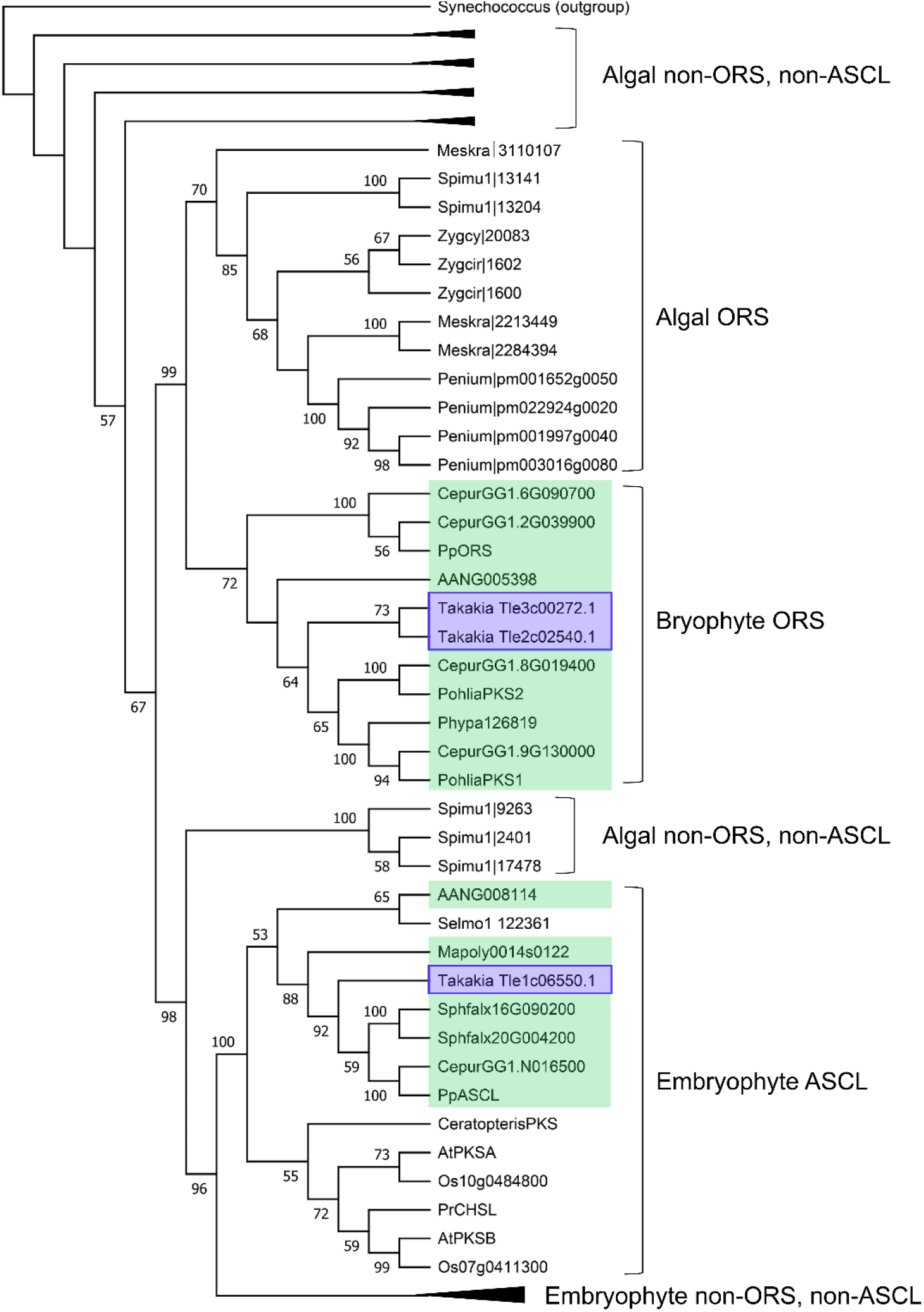
Maximum Likelihood tree of plant type III polyketide synthases. Included in the tree are sequences of all known algal type III PKS enzymes, all bryophyte enzymes that belong to either ORS or ASCL clades, representative ASCL enzymes from major embryophyte groups, and a selection of non-ORS and non-ASCL type III PKS enzymes from diverse embryophyte taxa. Bootstrap values (>50%) are displayed at the nodes. A cyanobacterial type III PKS was used as outgroup to root the tree. For brevity, four of the five algal non-ORS and non-ASCL clades and the embryophyte non-ORS and non-ASCl clade have been collapsed.

## Results

The ML phylogenetic tree (Fig. 1) resolved the type III PKS sequences into the following clades: **(a)** an embryophyte ASCL clade containing bryophyte ASCL sequences including one *Takakia* sequence plus a selection ASCL sequences from other major embryophyte groups. In agreement with the other ASCL sequences, the *Takakia* ASCL possesses diagnostic sequence features in addition to those for type III PKSs, namely Gly225 and (Ala/Val)240 (numbering based on PpASCL (Colpitts et al. 2011)), **(b)** a bryophyte clade containing exclusively ORS sequences including two *Takakia* sequences. In agreement with the other ORS sequences, the *Takakia* ORSs possess diagnostic sequence features in addition to those for type III PKSs, namely Gln218, (Val/Ala)277 and Ala286 (numbering according to PpORS (Kim et al. 2013)), **(c)** an algal clade containing exclusively Charophyte ORS sequences, **(d)** an embryophyte clade comprised of a selection of non-ORS and non-ASCL type III PKS sequences including a representative *Takakia* type III PKS, **(e)** five separate algal clades, collectively comprising 35 non-ORS and non-ASCL type III PKS sequences.

## Discussion

*Takakia lepidozioides* has a single *ASCL* gene in agreement with our contention that all embryophytes, with the possible exception of a few species, possess ASCL and therefore genuine SP in their spore or pollen walls. The few species predicted to lack ASCL are likely to exist in habitats that do not require protection from subaerial stresses, e.g. *Zostera marina*, and are presumed to have lost ASCL and pollen wall exine secondarily by reductive evolution.

We have reinforced our discovery that, while algae possess type III PKS sequences, none of them fall within the ASCL clade and, therefore, algae don’t possess genuine SP.

*Takakia* has four ORS sequences, two of which are truncated and consequently were omitted from our phylogenetic tree. We have shown previously that ORS is required for integrity of the leaf cuticle of *P. patens* and for its resistance to dehydration (Li et al. 2018) and that 2′-oxoalkylresorcinols restore dehydration tolerance in a PpORS knockout line (Aslam et al. 2022). We presume ORS has the same role in *Takakia* and other bryophytes. Interestingly, ORS sequences are present in Charophytes but appear to be absent from other algae. At present, there are no published reports about the location of ORS products in Charophytes or their roles.

